# Metaviromic identification of genetic hotspots of coronavirus pathogenicity using machine learning

**DOI:** 10.1101/2020.08.13.248575

**Authors:** Jonathan J. Park, Sidi Chen

## Abstract

The COVID-19 pandemic caused by SARS-CoV-2 has become a major threat across the globe. Here, we developed machine learning approaches to identify key pathogenic regions in coronavirus genomes. We trained and evaluated 7,562,625 models on 3,665 genomes including SARS-CoV-2, MERS-CoV, SARS-CoV and other coronaviruses of human and animal origins to return quantitative and biologically interpretable signatures at nucleotide and amino acid resolutions. We identified hotspots across the SARS-CoV-2 genome including previously unappreciated features in spike, RdRp and other proteins. Finally, we integrated pathogenicity genomic profiles with B cell and T cell epitope predictions for enrichment of sequence targets to help guide vaccine development. These results provide a systematic map of predicted pathogenicity in SARS-CoV-2 that incorporates sequence, structural and immunological features, providing an unbiased collection of genetic elements for functional studies. This metavirome-based framework can also be applied for rapid characterization of new coronavirus strains or emerging pathogenic viruses.

## Introduction

The coronavirus disease 2019 (COVID-19) pandemic caused by the severe acute respiratory syndrome coronavirus 2 (SARS-CoV-2) has become an unprecedented on-going global public health and economic crisis since its emergence at the end of the year 2019 ^1,2^. The SARS-CoV-2 virus has infected more than 5.4 million people and caused over 340,000 deaths globally as of May 25^th^, 2020 ^3^. Although pathogenic coronaviruses have repeatedly emerged from the wild to become infectious to human populations, the common genetic and molecular features that drive the disease-causing potential of these viruses are still unclear. Identifying genetic elements and specific regions of the SARS-CoV-2 genome that make it dangerous is critical for public health prevention and disease mitigation, as well as the development of vaccines and therapeutics. To address the current lack of systematic understanding of the pathogenic genomic features for coronaviruses, we developed a set of machine learning (ML) approaches focused on unbiased scanning and scoring of key pathogenicity-linked regions in the genomes of SARS-CoV-2 and other respiratory disease-causing coronaviruses ^4^ that distinguish them from non-pathogenic coronaviruses. We also harness these toolkits to identify genetic elements at high resolution that interface with critical viral protein functions and immunogenicity for vaccine development.

## Results

### Ensemble machine learning and statistical meta-model identifies high-resolution pathogenic features in coronavirus genomes

We developed a rigorous, integrative ML-based approach to identify regions that contribute to coronavirus pathogenicity, incorporating Random Forests (RF), Support Vector Machines (SVM), Bernoulli Naïve Bayes (BNB), Gradient Boosting Classifiers (GBC) and Multi-layer Perceptron Classifiers (MLPC) (**Methods**) (**Figure 1A**). First, we aligned 3,665 Coronaviridae family genomes obtained from the Virus Pathogen Database and Analysis Resource (ViPR) database ^5^ with diverse taxonomic and host features (**Figure 1B**). We chose a set of five different supervised learning algorithms that have robust performance and represent methodological diversity including ensembles of decision trees, Bayes’ theorem, and neural networks. Then, we took an ensemble learning approach, where outputs from the multiple classifiers are combined to give higher confidence predictions, to train and evaluate 7,562,625 advanced ML models. To set up our predictor classes, we considered several classification strategies to capture signatures associated with pathogenicity (**Figure 1C**). We hypothesized that sequence features that enable coronaviruses to jump from animal populations to humans (strategy A) and that distinguish SARS-CoV, MERS-CoV and SARS-CoV-2 from other coronaviruses (strategy B) to likely be important contributors to pathogenicity. We also considered features that specifically distinguish SARS-CoV, MERS-CoV and SARS-CoV-2 that infect human hosts (strategy C) from all other coronaviruses. After training and evaluating our base ML models on six bp sliding windows tiled across the alignment (100,835 windows), we integrated performance accuracy scores into a statistically rigorous meta-model based on minimum entropy windows (**Methods**) to obtain biologically interpretable nucleotide-level coronavirus pathogenicity (NT-COPA) scores for every nucleotide in the SARS-CoV-2 genome (**Figure 1D**).

**Figure 1.**
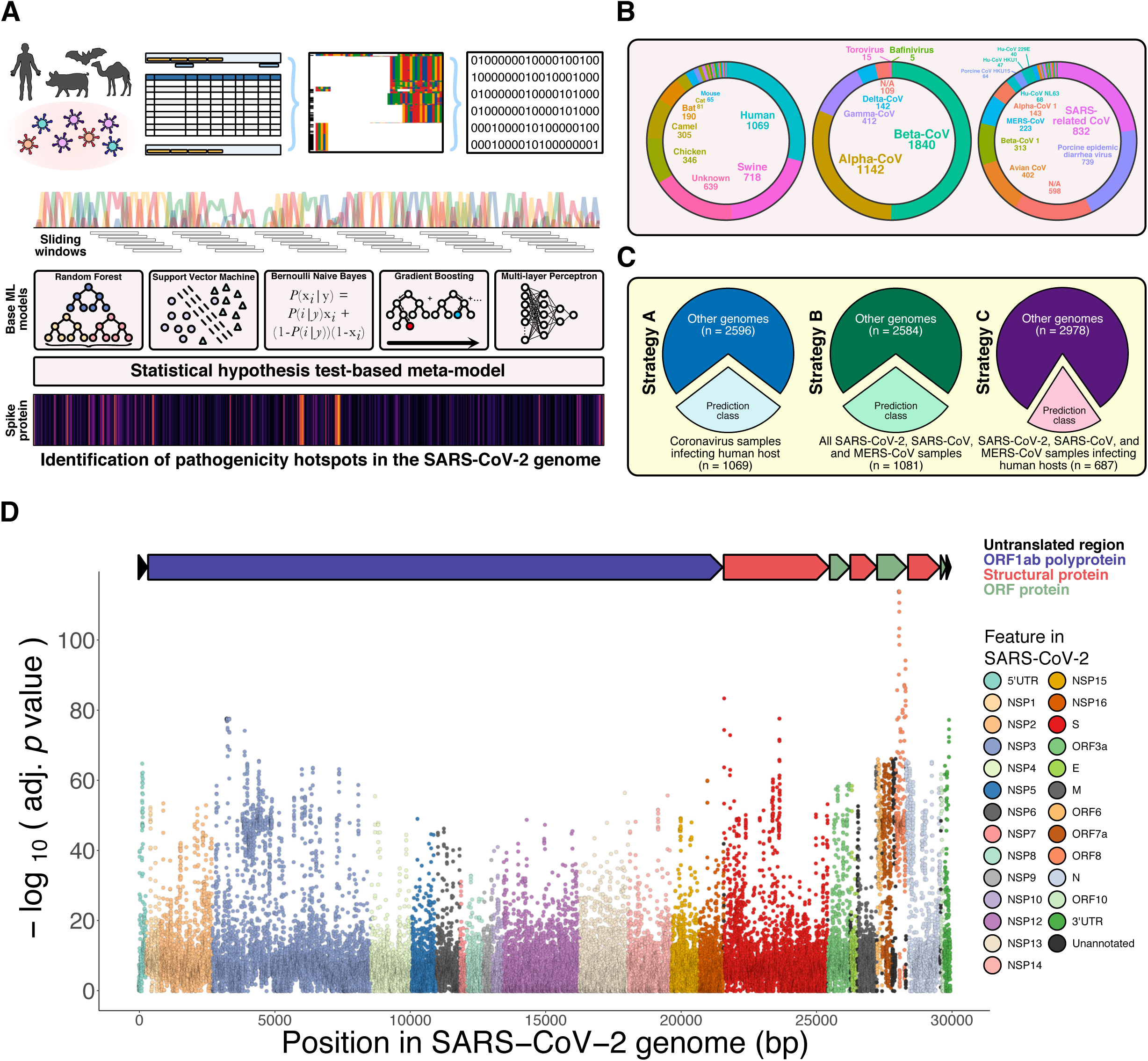
Ensemble machine learning and statistical meta-model identifies high-resolution pathogenic features in coronavirus genomes. **(A)** Schematic detailing ensemble machine learning strategy to learn pathogenic genomic features of coronaviruses. Complete genome sequences of the Coronaviridae family in the ViPR database (n = 3,665) were obtained, aligned, and encoded into binary vector representations. Base machine learning models with different classification strategies were trained on sliding windows tiled across the alignment. A statistical hypothesis test-based meta-model integrated signals into a coronavirus pathogenicity (COPA) score to identify pathogenicity hotspot regions in the SARS-CoV-2 genome. **(B)** (Left) Donut chart showing distribution of host species for coronavirus genomes used in study. (Middle) Donut chart showing the distribution of virus genus. (Right) Donut chart showing the distribution of virus species. **(C)** Pie charts showing class membership proportions for different classification strategies. (Left) Strategy A defines the predictor class as coronavirus samples that infect human hosts. (Middle) Strategy B defines the predictor class as all SARS-CoV-2, SARS-CoV, and MERS-CoV samples, including those that infect human or animal hosts. (Right) Strategy C defines the predictor class as specifically those SARS-CoV-2, SARS-CoV, and MERS-CoV samples that infect human hosts. **(D)** NT-COPA scores (negative log base 10 of adjusted p-values obtained from meta-model, see **Methods**) for every nucleotide position across the reference SARS-CoV-2 genome. Larger NT-COPA scores represent stronger pathogenicity signals learned from our models.

Empirically, ensemble approaches outperform single predictors and tend to yield better results when there is significant diversity in models ^6,7^. In addition to the three classification strategies, our base ML models covered five cross-validation folds (using 80% of samples for model training and 20% of samples for validation for a given permutation) and five different machine learning methods comprising SVC, RF, BNB, MLPC, and GBC. Because the samples in the standard dataset are ordered by alignment, individual models for different cross-validation folds may have dissimilar training compositions and therefore accuracy scores (**Figure S2**); however, this tradeoff may come with a greater diversity of biologically meaningful learned features. For comprehensiveness, we also trained our models on a dataset where genome samples are shuffled (**Figure S3**). While individual classifier performance increased, they also become more correlated (reducing model diversity) and led to higher saturation of NT-COPA score signals (**Figure S3D**) compared to results obtained from our standard dataset (**Figure 1D**). Since we pool together all cross-validation scores with training coverage of all samples, no genomic information should be lost for our statistical meta-model analyses with either method. We focused our subsequent analyses on results obtained from our standard dataset for biological interpretability, but provide NT-COPA scores obtained from both methods as a resource.

### Identification of local pathogenic hotspots in SARS-CoV-2 proteins

Next, we looked at NT-COPA score distributions intersected with the annotated SARS-CoV-2 genome to see if they can be used to identify potential pathogenicity hotspots. We found that the NT-COPA scores reflected quantitative and high-resolution signatures for characterizing individual base pairs and amino acids within SARS-CoV-2 features (**Figure 2A**). To address the challenge of systematically defining hotspot regions from such high-resolution data, we considered that scores for a given base pair should reflect local genomic information capture due to our sliding window based approach for training the base ML models. Therefore, we considered kernel smoothing to be an appropriate nonparametric curve estimation method for a region-based approach to identify hotspots (**Methods**). We calculated the kernel regression estimate at each base pair using the NT-COPA scores, and used the estimates to determine local signal maxima (peaks) within SARS-CoV-2 features. This approach yielded 2,473 peaks across the SARS-CoV-2 genome, which mark local pathogenicity hotspots (**Figure 2B**). Limitations to the kernel smoothing based approach include that identified peaks may have low NT-COPA scores as they only reflect local maxima (which may be addressed by using score thresholds), and that high signal density regions may only return a single peak. The advantage of the approach is that it is unbiased and systematic. Both systematic and customized strategies can be applied to generate biologically meaningful insights from these pathogenicity-associated scores.

**Figure 2.**
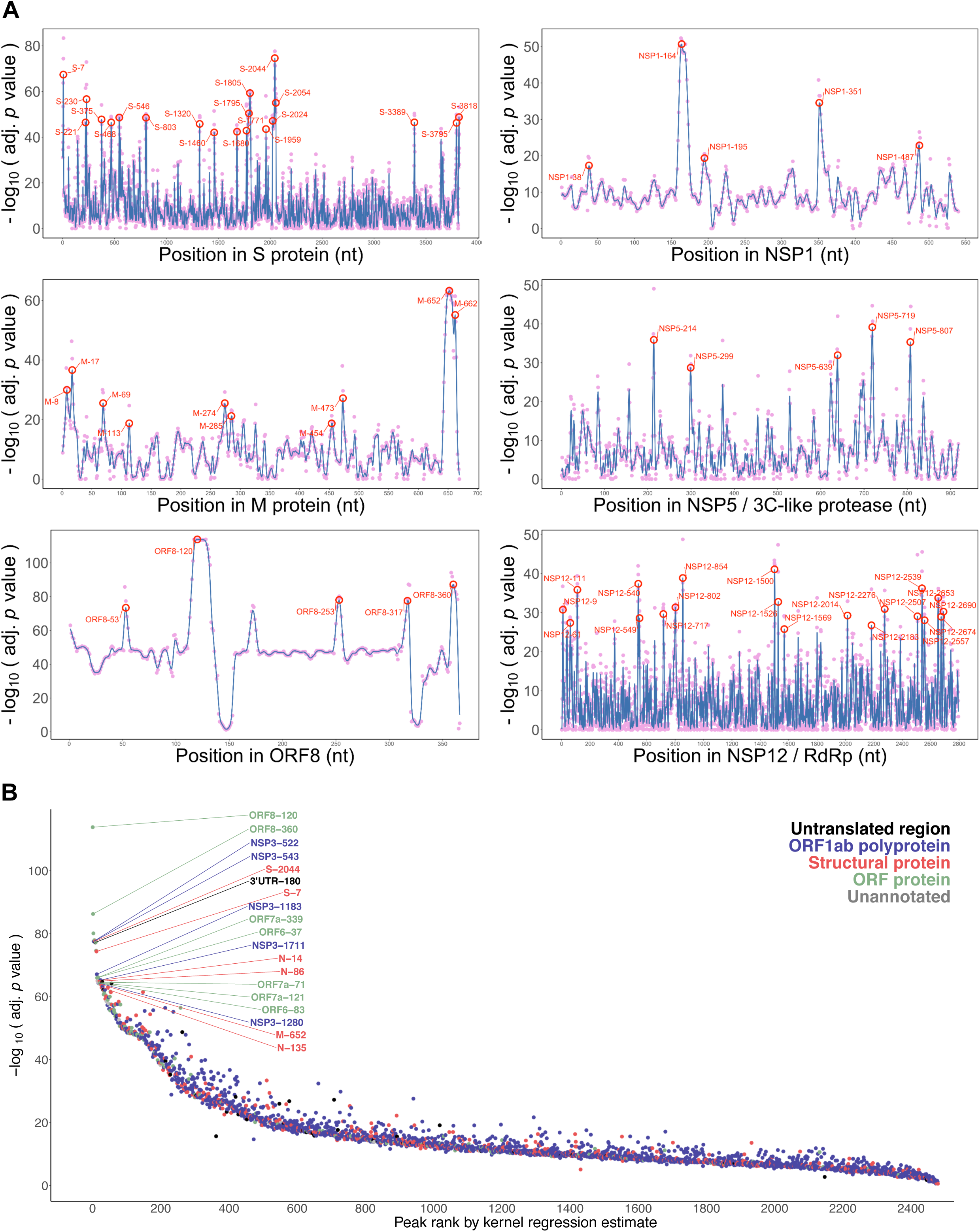
Identification of local pathogenic hotspots in SARS-CoV-2 proteins. **(A)** High resolution NT-COPA score distributions shown for spike protein, membrane protein, ORF8, NSP1, NSP5 (3C-like protease), and NSP12 (RNA-dependent RNA polymerase). Scores at each nucleotide position are shown as pink dots. Smoothed kernel regression estimates are shown a blue line graph, with select local peaks circled and labeled in red. NT-COPA scores for local pathogenicity-associated peaks identified across SARS-CoV-2 genome shown plotted against rank by kernel regression estimate. Smoothing and peak identification was used as an unbiased strategy to identify pathogenicity hotspots.

### Spike protein pathogenic hotspots reveal a furin cleavage site and contact sites with ACE2

To biologically validate the significance of our candidate hotspots, we performed a series of in-depth evolutionary and structural analyses. There has been considerable focus on the spike protein as it facilitates coronavirus entry into target cells ^8,9^. For SARS-CoV-2, interaction between the trimeric spike glycoprotein and the human host ACE2 receptor triggers a cascade of events that leads to the fusion between cell and viral membranes ^10^. We examined the NT-COPA score distributions and peaks for the spike protein and found the strongest signal to be peak S-2044, corresponding to amino acid position 682 (**Figure 2A**). In order to determine the evolutionary significance of this hotspot, we aligned the spike protein amino acid sequences for Coronaviridae family viruses across various species and hosts and compared the alignment with the NT-COPA score density for peak S-2044 and nearby residues (**Figure 3B**).

**Figure 3.**
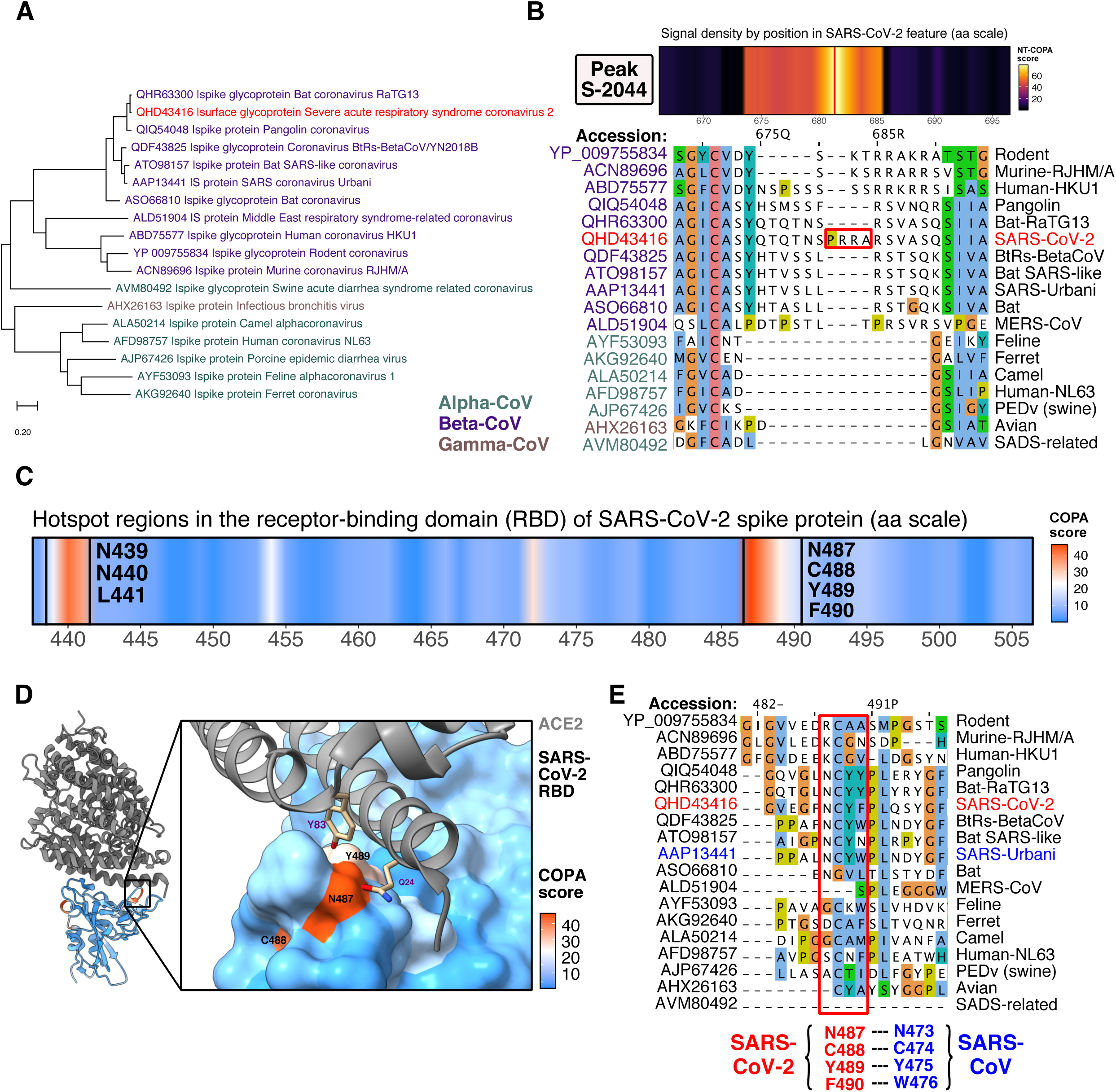
Spike protein pathogenic hotspots reveal a furin cleavage site and contact sites with ACE2. **(A)** Phylogeny tree for spike protein sequences of coronaviruses across species and hosts. Sequences for alphacoronavirus are labelled in green, betacoronaviruses labelled in purple, gammacoronaviruses labelled in brown, and the reference SARS-CoV-2 labelled in red. **(B)** NT-COPA score signal density near peak S-2044 (amino acid position 682) in spike protein compared to alignment. Peak S-2044 corresponds to a furin-like cleavage site. **(C)** AA-COPA score, i.e. COPA score signal density in receptor-binding domain (RBD) in spike protein reveals two primary pathogenicity hotspot regions. **(D)** COPA scores mapped onto structure of SARS-CoV-2 RBD complexed with ACE2 receptor reveal that the NCYF hotspot contains residues that mediate viral binding to host receptor. **(E)** Protein alignment reveals NCYF hotspot for SARS-CoV-2 (NCYW hotspot for SARS-CoV) has high sequence divergence from other coronaviruses across species and hosts. Residues are colored using the Clustal X color scheme. Hotspot residues for SARS-CoV-2 labelled in red, with corresponding residues for SARS-CoV labelled in blue.

We find that this peak corresponds to a functional polybasic furin cleavage site (RRAR) at the junction between the S1/S2 subunits, which has been reported to expand SARS-CoV-2 tropism and/or enhance its infectivity ^11^. The leading proline that is also inserted at the site for SARS-CoV-2 (for PRRA insertion) has been shown to result in addition of O-linked glycans to S673, T678 and S686 which flank the cleavage site by structural analysis ^12^. Nucleotides for the T678 codon and the first nucleotide for the S686 codon (corresponding to position S-2056) all have high NT-COPA scores and are included as part of the peak S-2044 associated hotspot. More generally, this hotspot, which spans nucleotide positions 2,021 to 2,056 (amino acid positions 674 to 686, all with NT-COPA scores > 40), corresponds to amino acid insertions that contribute to distinguishing betacoronaviruses from alpha- and gammacoronaviruses (**Figure 3A-B**). The functional consequence of the polybasic cleavage site and the predicted O-linked glycans in SARS-CoV-2 remains unclear, although possibilities for the latter include creation of mucin-like glycan shields involved in immune evasion ^12^. These analyses showed that the ML-based approach independently learned pathogenicity signals that correspond to important features of the SARS-CoV-2 genome, several of which have been previously validated or are under active investigation.

To see if ML-scored pathogenicity hotspots can offer functionally significant structural insights, we examined the spike protein receptor-binding domain (RBD) interface with ACE2. We calculated the amino acid resolution COPA scores (AA-COPA, or COPA for short) by averaging the NT-COPA scores for codons. We then examined the high AA-COPA regions in the spike protein RBD, and identified two hotspot regions comprising residues NNL at positions 439-441 and residues NCYF at positions 487-490 (**Figure 3C**). We then mapped the COPA scores onto a recently solved crystal structure of the wild-type SARS-CoV-2 RBD bound to human ACE2 ^10^, and found that the NCYF hotspot included contact site residues at the RBD-ACE2 interface (**Figure 3D**). Of the 13 hydrogen bonds at the SARS-CoV-2 RBD - ACE2 interface identified from the wild-type structure (**Figure S5B**), 3 hydrogen bonds are included in the NCYF hotspot: N487-Q24, N487-Y83, and Y489-Y83. Notably, all three of these SARS-CoV-2 - ACE2 hydrogen bonds are conserved for the SARS-CoV RBD - ACE2 interface, as N473-Q24, N473-Q24, Y475-Y83 ^10^. Both of the coronavirus contact site residues in the SARS-CoV-2 NCYF hotspot (N487 and Y489) are relatively conserved amongst proximal strains, but differ in less proximal strains (**Figure 3A, 3E**), suggesting that acquisition of these sites were important evolutionary events in development of high affinity coronavirus binding to the human ACE2 receptor. Interestingly, an alternative, chimeric RBD-engineered structure of the SARS-CoV-2 spike protein-ACE2 complex demonstrated that structural changes in one of the ridge loops that differentiate SARS-CoV-2 from SARS-CoV introduces an additional main-chain hydrogen bond between residues N487 and A475 in the SARS-CoV-2 receptor binding motif (RBM), causing the ridge to form more contacts with the N-terminal helix of ACE2 ^13^. The COPA-structural joint analysis suggested that the ML models automatically learned the SARS-CoV-2 NCYF hotspot as a proximally conserved contributor to coronavirus pathogenicity.

We then examined the other hotspot region identified in the SARS-CoV-2 RBD, comprising residues N439, N440, and L441. Residue N439 was not identified to be involved in contacts between SARS-CoV-2 RBD and ACE2 receptor in the wild-type structure (**Figure S5C**). However, its associated residue in the SARS-CoV RBD, R426, forms a strong salt bridge with E329 on ACE2 and a hydrogen bond with Q325 ^10,13,14^ (**Figure S5D**). Evolutionary analysis reveals that the NNL (SARS-CoV-2 coordinates) or RNI (SARS-CoV coordinates) hotspot has substantial sequence divergence from other coronaviruses across species and hosts (**Figure S5A**). While the significance for the NNL hotspot for SARS-CoV-2 is unclear, R426 is a functionally important residue for ACE2 receptor binding in SARS-CoV, and scored highly in the classification strategies focused on learning sequence determinants of pathogenicity that are generalizable across respiratory disease-causing coronaviruses.

### RdRp pathogenic hotspots reveal RNA contact sites and codon composition biases

Another key component of the SARS-CoV-2 virus is the RNA-dependent RNA polymerase (RdRp), also known as nonstructural protein 12 (NSP12). RdRp/NSP12 forms a complex with accessory factors including NSP7 and NSP8, which increase template binding and processivity, to catalyze the synthesis of viral RNA ^15,16^. As this complex plays an important role in the viral replication and transcription cycle, RdRp is currently being investigated as a target of nucleotide analog antiviral drugs such as remdesivir for COVID-19 treatment ^17,18^. To identify hotspot regions in RdRp associated with pathogenicity, we intersected its sequence with the ML-generated COPA scores (**Figure 4A**). We then mapped the COPA scores onto a recently solved cryo-EM structure of the SARS-CoV-2 NSP12-NSP7-NSP8 complex bound to template-primer RNA and the triphosphate form of remdesivir (RTP) ^16^ (**Figure 4B**). We focus on two structural regions of interest in SARS-CoV-2 RdRp with high COPA score signal density. Region (1), which comprises residues ERVRQ (positions 180-184) and DRY (positions 284-286), reflects a previously uncharacterized feature of RdRp with a high density of hydrophobic and hydrophilic amino acid residues. Whether the hotspot residues in region (1) create networks of hydrophilic interactions that contribute to pathogenicity require further experimental study; nevertheless, this region highlights pathogenicity features that were learned from the ML models in an unbiased manner. Region (2), which comprise residues K500, S501, W509, and I847, includes key residues involved in direct RdRp protein-RNA interactions. We observed that the identified COPA hotspot residues generally exhibit high amino acid conservation amongst proximal strains and differentiation in less proximal strains (**Figure 4C and Figure S6B**), with a notable exception of residues K500 and S501.

**Figure 4.**
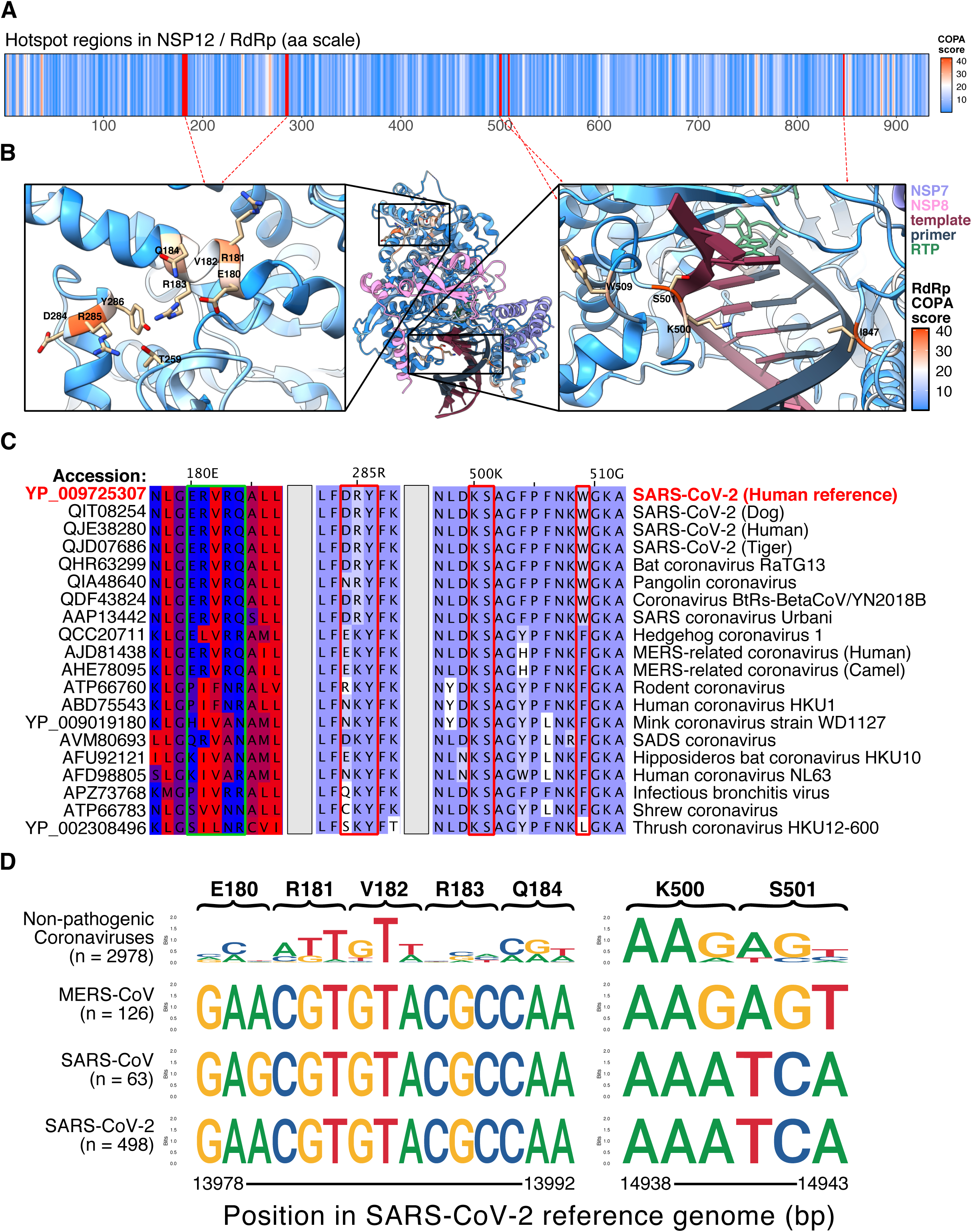
RdRp pathogenic hotspots reveal RNA contact sites and codon composition biases. **(A)** COPA score signal density across the NSP12/RdRp amino acid sequence. Select hotspot residues marked in red and directed towards their location on structure in **(B)**. **(B)** COPA scores mapped onto RdRp in structure of SARS-CoV-2 RdRp-NSP7-NSP8 complex bound to the template-primer RNA and triphosphate form of remdesivir (RTP). (Left) Spatial region in RdRp with high density of pathogenicity hotspots. (Right) Pathogenicity hotspot residues correspond to contact sites in RdRp that directly participate in the binding of RNA. **(C)** Protein alignment reveals pathogenicity hotspot residues in SARS-CoV-2 RdRp has high sequence divergence from other coronaviruses across species and hosts, with exceptions as noted in **(D)**. For the ERVRQ hotspot region (amino acid positions 180 to 184), residues are colored according to hydrophobicity (where most hydrophobic residues are colored red and the most hydrophilic ones are colored blue). For other regions, residues are colored according to their Blosum62 score (where residues matching the consensus sequence residue at that position are colored dark blue). **(D)** Sequence logos for codons from the genome alignment used for ML training associated with the ERVRQ hotspot region and KS hotspot contact sites. Logos were generated separately for MERS-CoV, SARS-CoV, and SARS-CoV-2 infecting human hosts, and non-pathogenic coronaviruses.

We were surprised at this exception since initially it was unclear why our ML approach would assign residues K500 and S501 high COPA scores if these positions exhibit such strong evolutionary conservation across species and hosts, as these positions should then not be able to distinguish pathogenic coronaviruses. To examine these regions at the nucleotide resolution, we returned to our aligned genome used for training the base ML models, and generated nucleotide composition frequencies (presented as motifs of sequence logos) for codons associated with the hotspot residues (**Figure 4D)**. The ERVRQ motif reveals conservation amongst the pathogenic coronaviruses that differentiate them from the high diversity of non-pathogenic coronaviruses in this region. These results are expected given the goals of our methods. The KS motif, however, reveals codon composition bias that differentiate SARS-CoV-2 and SARS-CoV at the nucleotide level from MERS-CoV and non-pathogenic coronaviruses. This bias is particularly striking for residue S501, where all three nucleotides differentiate the SARS strains from other coronaviruses, despite conserving a serine residue. Whether these codon composition biases reflect selection, recombination, or more generalized codon usage biases require further study. Nevertheless, these results highlight the learned evolution signatures of critical features in the SARS-CoV-2 genome at different levels.

### Integration of genomic pathogenicity profiles with B cell and T cell immunogenic features

There is a dire need for rapid development of effective vaccines to fight the COVID-19 pandemic, but given the novelty of the virus there are currently none approved specifically for SARS-CoV-2. Although a few vaccine candidates are in active clinical trials (e.g. Moderna, Pfizer/BioNtech), most of the current approaches use the spike glycoprotein as a target and primarily use full length or simple partial ORFs ^19^. It is unclear if this single target will prove to be sufficient for mounting protective and prolonged immunity for humans. More generally, limited information on which parts of the virus are recognized by human immune responses is a major knowledge gap impeding effective vaccine design, although efforts are currently underway to study patterns of immunodominance ^20^ and to identify conserved epitopes for cross-reactive antibody binding ^21^. While current vaccine strategies focus on inducing B cell humoral responses, T cell immunity comprises another dominant domain of immune responses essential for viral vaccines ^22–24^ and may play an important role in eliminating SARS-CoV-2 ^20,25^. Therefore, it is important to examine both B cell and T cell epitopes and consider more precise pathogenic and immunogenic regions that could potentially induce a stronger immune response.

We thus set out to identify those regions by intersecting both pathogenic hotspots and immunogenic hotspots. To identify regions in the SARS-CoV-2 proteome that are predicted to be both pathogenic and immunologically relevant, we ran B cell epitope analysis (**Figure 5A**) as well as T cell MHC-I and MHC-II binder predictions, and then integrated them with the ML-generated COPA scores (**Figure 5B, Figure 6B, and Figure 7A**). Surprisingly, we found that for spike and nucleocapsid proteins, high COPA pathogenic regions significantly overlap with potential B cell epitopes (hypergeometric test, p < 0.008 for spike, and p < 0.0012 for nucleocapsid) (**Figure 5A, Figure 6A**). For T cell epitopes, we prioritize peptides by counts of pathogenic hotspot peaks obtained from the kernel smoothing analysis. These convergent regions may enrich for highly immune-reactive elements and help epitope prioritization to guide vaccine target selection more precisely on functionally important regions of SARS-CoV-2 (**Figure 5B, Figure 6B, Figure 7A**). In addition to potentially mounting stronger immunity, incorporating pathogenicity signals may help developing vaccines that generate immune responses enriched in neutralization of the more dangerous viral elements. We join other efforts for systematic characterization of SARS-CoV-2 features ^26^and provide in this study all the regional hotspots as consensus regions for next-generation precision vaccine development.

**Figure 5.**
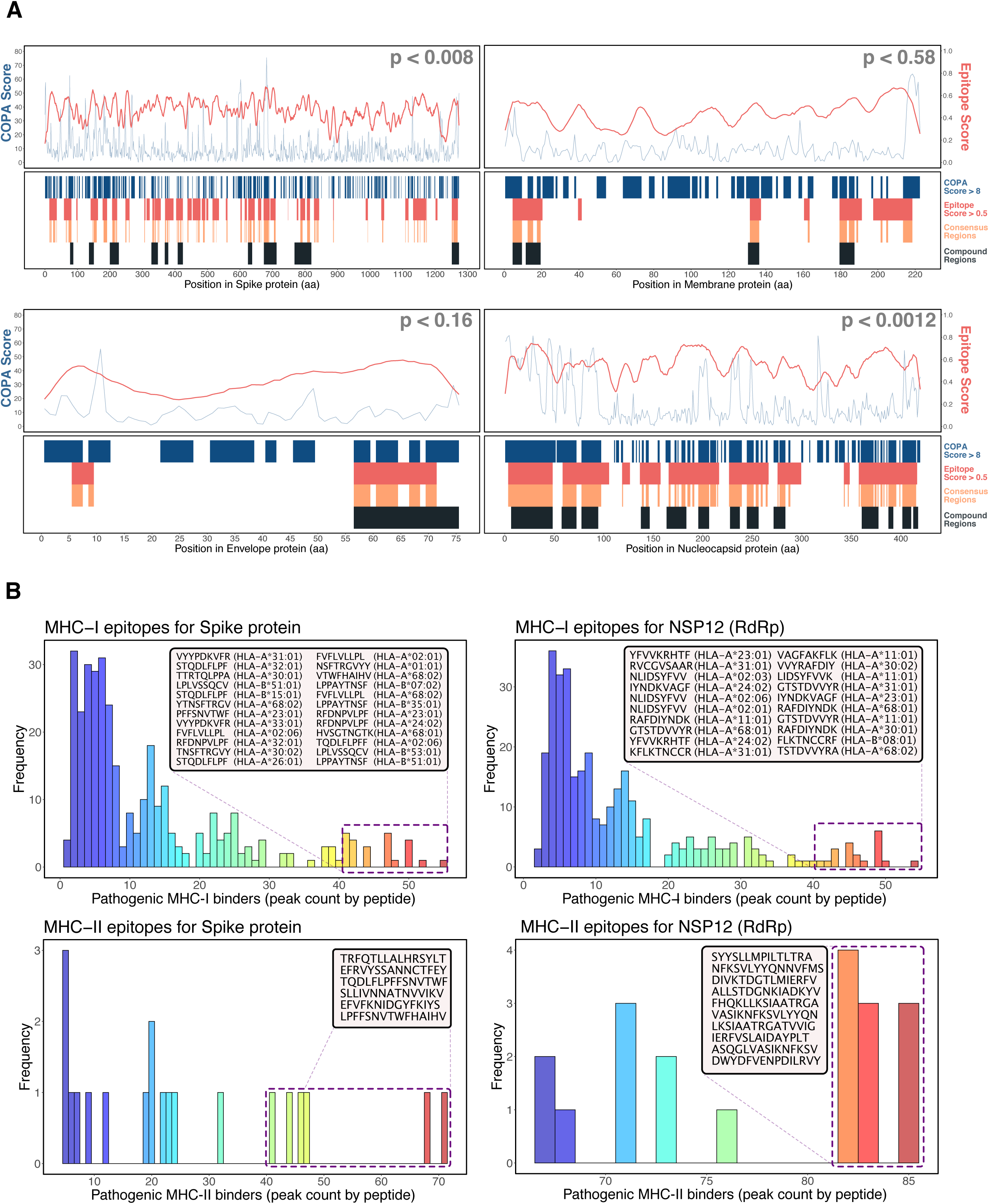
Integration of genomic pathogenicity profiles with B cell and T cell immunogenic features. **(A)** B cell epitope integrative analyses for spike protein, membrane protein, envelope protein, and nucleocapsid protein. (Upper) COPA scores and B cell epitope prediction scores plotted across the amino acid sequences. Thresholds used for identifying key residues (COPA score > 8 and epitope score > 0.5) marked with horizontal line. Statistical significance of overlap of key residues was determined by hypergeometric test (see Figure 6A). (Lower) Residues marked for COPA score > 8, epitope score > 0.5, consensus regions, and compound regions. **(B)** T cell epitope integrative analyses for spike protein and RdRp for MHC-I and MHC-II. Pathogenicity-associated peaks identified from kernel regression estimates were mapped onto predicted MHC-I and MHC-II binders. Peptides with high peak counts are highlighted.

**Figure 6.**
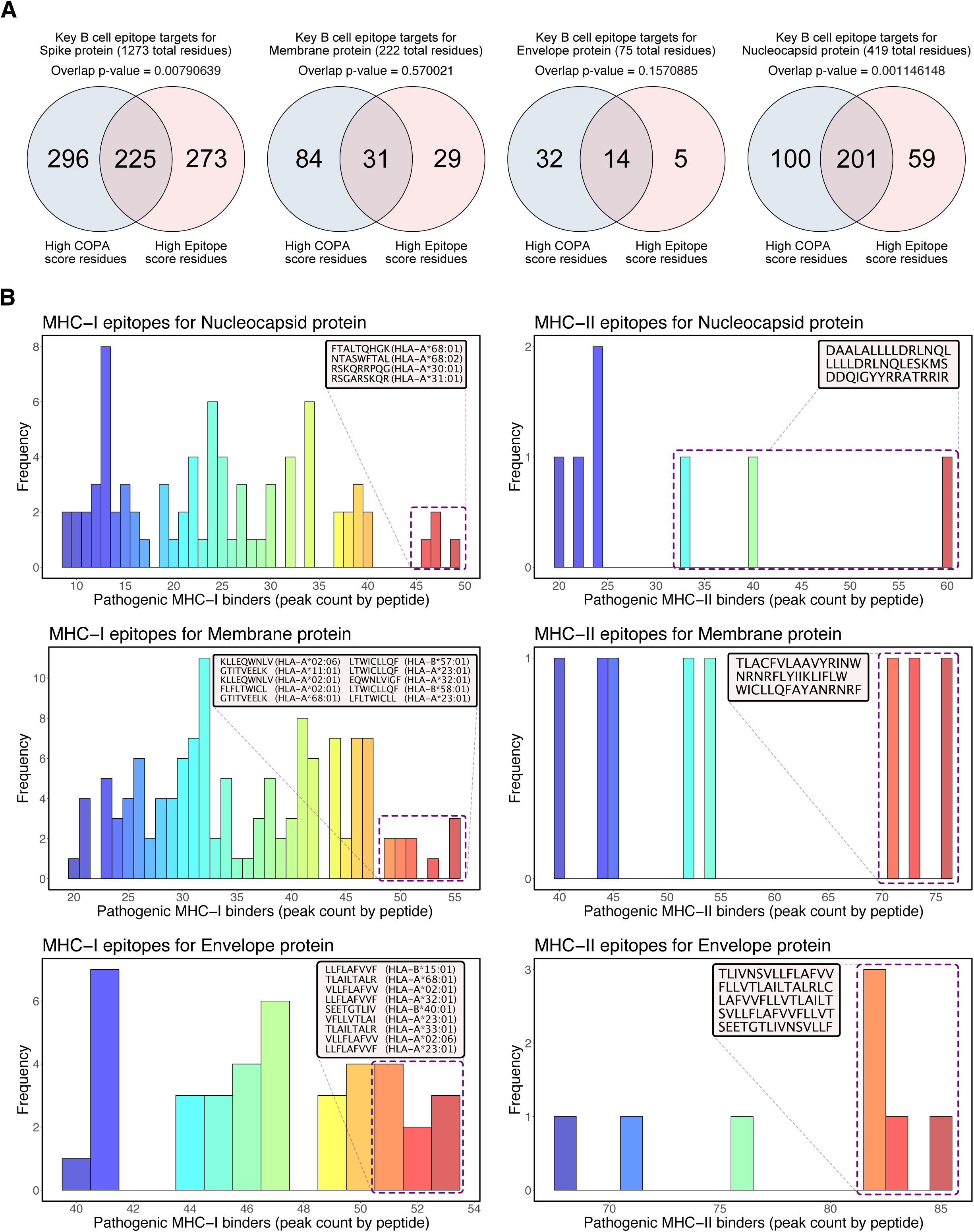
Additional integrative analyses of pathogenicity profiles with B cell and T cell epitopes for SARS-CoV-2 structural proteins. **(A)** Venn diagrams showing overlap of key residues in spike protein, membrane protein, envelope protein, and nucleocapsid protein identified with thresholds of COPA score > 8 and epitope score > 0.5. Statistical significance of overlap of key residues was determined by hypergeometric test. **(B)** T cell epitope integrative analyses for nucleocapsid protein, membrane protein, and envelope protein for MHC-I and MHC-II. Pathogenicity-associated peaks identified from kernel regression estimates were mapped onto predicted MHC-I and MHC-II binders. Peptides with high peak counts are highlighted.

**Figure 7.**
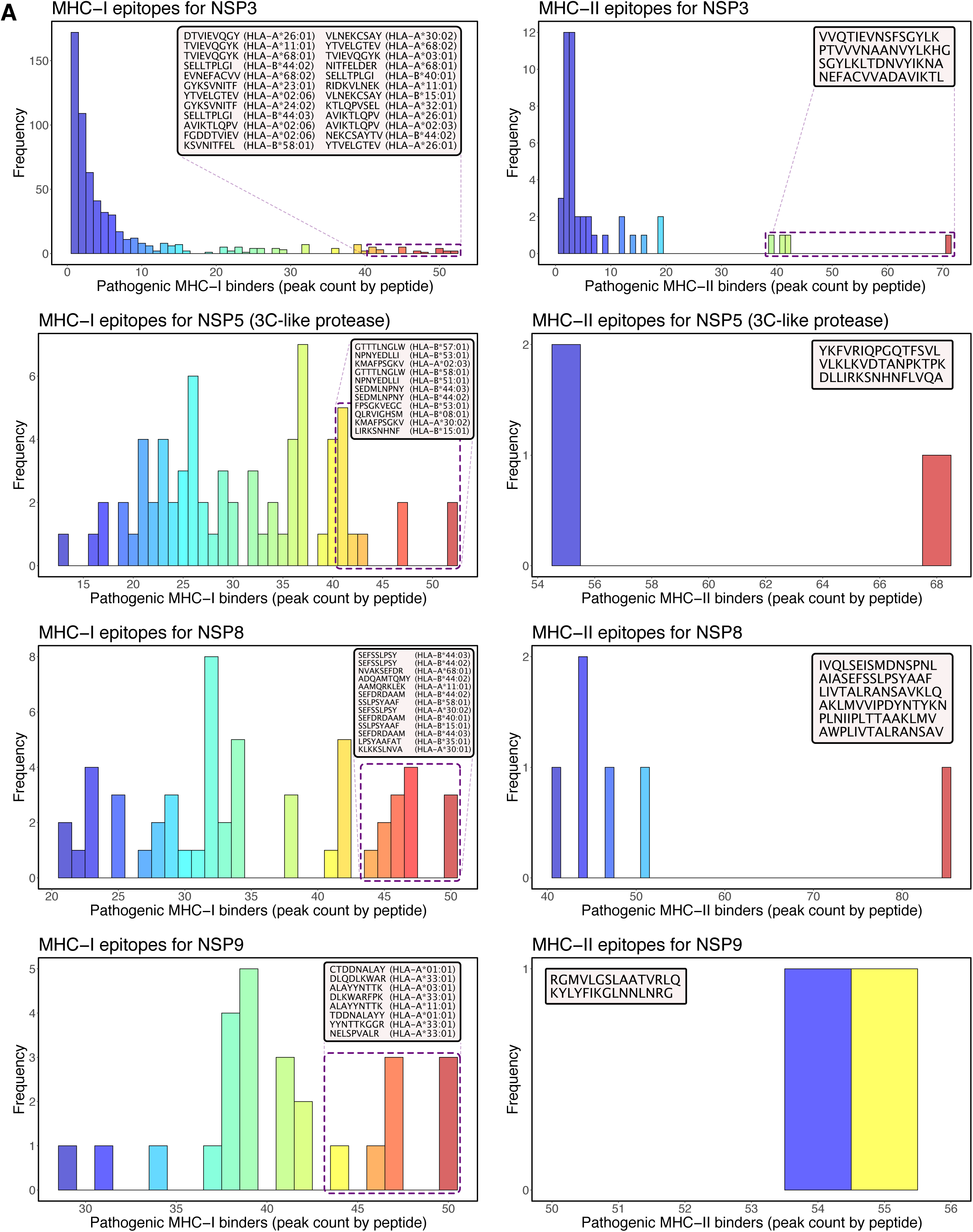
Additional integrative analyses of pathogenicity profiles with T cell epitopes for SARS-CoV-2 nonstructural proteins. **(A)** T cell epitope integrative analyses for NSP3, NSP5, NSP8, and NSP9 for MHC-I and MHC-II. Pathogenicity-associated peaks identified from kernel regression estimates were mapped onto predicted MHC-I and MHC-II binders. Peptides with high peak counts are highlighted.

## Discussion

This study developed a rigorous ensemble learning framework that integrates base ML models and a statistical meta-model to distinguish pathogenic sequence features of coronaviruses down to base pair and amino acid resolutions with quantitative and biologically interpretable COPA scores. By training and evaluating a high number of diverse ML models on a large collection of coronavirus genomes of human and animal origins, we identified pathogenic hotspots across the SARS-CoV-2 viral genome. Comparative validation with previous work through in-depth, biologically-motivated investigation showed various intersections of common key features, while the ML approach itself is fully unbiased in terms of scoring and generation of a large number of previously unidentified candidate hotspots. For example, the significance of these hotspots were shown with in-depth evolutionary and structural analyses of the spike protein and RdRp, which are important SARS-CoV-2 genetic elements under active investigation. The integrative analysis of pathogenicity-associated genomic profiles with B cell and T cell epitopes converged on regions of the SARS-CoV-2 proteome that are predicted to be both pathogenic and immunologically relevant, which provides a collection of feature-rich elements that potentially serve as candidates for prioritization and enrichment of key sequence targets to guide vaccine development.

While we focused our downstream analysis here on spike protein and RdRp to demonstrate the interpretability and functional significance of the COPA scores learned from our framework, we emphasize that the learned features from this study are genome-wide and may provide insights into less characterized SARS-CoV-2 structural and non-structural proteins. For example, we noticed that our framework identified a high density of pathogenic peaks in ORF8 (**Figure 2B**), a protein whose function remains mysterious ^27^. A recent pre-print has identified a 382-nt deletion variant that covers nearly the entirety of ORF8 from strains isolated from hospitalized patients in Singapore ^28^, which were implied to lead to reduced virulence of SARS-CoV-2 based on experimental data from SARS-CoV OFR8 deletion variants ^29^. Whether ORF8 is in fact an important driver of SARS-CoV-2 pathogenicity will require further study; meanwhile, the pathogenicity signatures identified here may constitute an unbiased collection for regional dissection through viral experiments.

Given the ongoing nature of the COVID-19 pandemic, there is an urgent need to identify functionally important features of SARS-CoV-2. While much effort is currently underway to characterize the spike protein, RdRp, and other proteins suggested to be important from prior studies on coronaviruses, there has been limited information on sequence determinants of pathogenicity at the global, metavirome-wide scale. We demonstrate here how harnessing the predictive power of ML or other artificial intelligence algorithms may be used to identify such features in a systematic manner. While our ML strategies are based on primary sequences, future ML algorithms that incorporate 3D structures may generate additional insights that cannot be obtained from linear sequence analysis alone, and further enhance the prediction of pathogenicity, immunogenicity, or other important elements of viral proteins. This study demonstrates the development and application of ML to coronavirus genomes with integrative analyses, which is not limited to coronaviruses but can be broadly applied to other viral genomes or microbial pathogens to gain insights on pathogenicity and immunogenicity.

## Acknowledgments

We thank Richard Sutton, Hongyu Zhao, Albert Ko, Yong Xiong, Craig Wilen, Akiko Iwasaki, Katie Zhu, Ruth Montgomery, and a number of other colleagues for discussions on COVID-19. We thank Antonio Giraldez, Andre Levchenko, Chris Incarvito, Mike Crair, and Scott Strobel for their support on COVID-19 research. We thank Chen lab members such as Matthew Dong, Vino Peng, Ryan Chow, Paul Clark, Stanley Lam, as well as our colleagues in the Genetics Department, the Systems Biology Institute and various Yale entities. We personally thank all the frontline healthcare workers directly fighting this disease.

## Author contributions

JJP and SC conceived and designed the study. JJP developed the analysis approach, performed all data analyses, and created the figures. JJP and SC prepared the manuscript. SC supervised the work.

## Declaration of interests

No competing interests related to this study.

The authors have no competing interests, or other interests that might be perceived to influence the interpretation of the article. The authors are committed to freely share all COVID-19 related data, knowledge and resources to the community to facilitate the development of new treatments or prevention approaches against SARS-CoV-2 / COVID-19 as soon as possible.

As a note for full disclosure, SC is a co-founder, funding recipient and scientific advisor of EvolveImmune Therapeutics, which is not related to this study.

## Methods

### Sequence data collection

A total of 3,665 complete nucleotide genomes of the “Coronaviridae” family were downloaded from the Virus Pathogen Database and Analysis Resource (ViPR) database ^5^ to be used for machine learning algorithm training. Genbank accession MN908947 was used as the reference SARS-CoV-2 sequence for downstream analyses. Coronavirus protein sequences for spike protein (YP_009755834, ACN89696, ABD75577, QIQ54048, QHR63300, QHD43416, QDF43825, ATO98157, AAP13441, ASO66810, ALD51904, AYF53093, AKG92640, ALA50214, AFD98757, AJP67426, AHX26163, AVM80492) and ORF1ab (QIT08254, QJE38280, QJD07686, QHR63299, QIA48640, QDF43824, AAP13442, QCC20711, AJD81438, AHE78095, ATP66760, ABD75543, YP_009019180, AVM80693, AFU92121, AFD98805, APZ73768, ATP66783, YP_002308496) used for evolutionary analyses were obtained from the NCBI Virus community portal. Amino acid sequences for SARS-CoV-2 were obtained from translations from reference sequence NC_045512 (equivalent to MN908947). FASTA sequences for S protein (YP_009724390), E protein (YP_009724392), M protein (YP_009724393), N protein (YP_009724397), NSP3 (YP_009742610), NSP5 (YP_009742612), NSP8 (YP_009742615), NSP9 (YP_009742616), and NSP12 (YP_009725307) were obtained from the NCBI Protein database and used for downstream evolutionary and immune epitope analyses.

### Pre-processing

Sequences were aligned with MAFFT ^30^ version 7 with the --auto strategy. Degenerate IUPAC base symbols that represent multiple bases were converted to “N” and ultimately masked prior to training algorithms. Six bp-wide sliding windows with 1bp shifts were generated across every position in the alignment for a total of 100,835 alignment-tiled windows. Genetic features including nucleotides and gaps for a given window were converted to binary vector representations using LabelEncoder and OneHotEncoder from the Python scikit-learn library ^31^, for integer encoding of labels and one-hot encoding respectively. Additional Python libraries used include BioPython ^32^, NumPy ^33^, and pandas ^34^.

### Training and evaluating machine learning base models

Genome metadata was converted to binary vector classifications with “1” representing predictor class genomes depending on classification strategy and “0” representing all other genomes. Three different classification strategies were used: (1) predictor class comprised coronavirus samples infecting human hosts, (2) predictor class comprised all SARS-CoV-2, SARS-CoV, and MERS-CoV samples, and (3) predictor class comprised SARS-CoV-2, SARS-CoV, and MERS-CoV samples specifically infecting human hosts. Five supervised learning classifiers from scikit-learn were used for training and evaluation, with seeds set at 17 for algorithms that use a random number generator. Support vector classifiers (SVC) were trained with a linear kernel and regularization parameter of 1.0; random forest (RF) classifiers were trained with 100 estimators; Bernoulli Naïve Bayes (BNB) were trained with alpha of 1.0 with the “fit_prior” parameter set as true to learn class prior probabilities; multi-layer perceptron (MLPC) classifiers were trained with “lbfgs” solver, alpha of 1e-5, 5 neurons in the first hidden layer, and 2 neurons in the second hidden layer; gradient boosting classifiers (GBC) were trained with “deviance” loss function, learning rate of 0.1, and 100 estimators. All estimators were trained and evaluated with stratified 5-fold cross-validation on each window, using 80% of the data for training and 20% of the data for validation.

### Statistical hypothesis test-based meta-model

Accuracy scores obtained from machine learning base models were used as a proxy for “learned, predictive information content” to determine coronavirus pathogenicity (COPA) scores using a statistical hypothesis test-based meta-model. First, Shannon entropy values were calculated for each window across the alignment. Windows with minimal entropy values (n = 10,383), typically found in highly gapped regions, were used to define a biologically meaningful control group; i.e., we hypothesized that windows with low information content in highly gapped regions should not be predictive of coronavirus pathogenicity. For each position across the alignment (100,840 positions), scores associated with windows that overlap with the position (typically ∼six windows) were pooled and tested to see if statistically significantly different from the minimal entropy control group using the nonparametric two-sided Wilcoxon rank-sum test. This procedure was performed across the alignment, and p-values were adjusted for multiple comparisons using the Benjamini & Hochberg procedure. P-values were transformed to nucleotide resolution coronavirus pathogenicity scores by negative log base 10 (also referred to as NT-COPA scores). Amino acid resolution scores were obtained by averaging the NT-COPA scores for a given residue’s codon (referred to simply as COPA scores).

### Kernel regression smoothing for hotspot peak identification

For a systematic strategy to identify pathogenicity hotspots across the SARS-CoV-2 genome using COPA scores, we combined kernel regression smoothing with local maxima identification. For each position across the alignment, we determined the Nadaraya-Watson kernel regression estimate using the ksmooth function in R with a “normal” kernel and various bandwidth sizes. Peaks highlighted in this study are primarily based on estimates calculated with bandwidth size of 3. Local peaks were determined from kernel regression estimates using the “findpeaks” function with nups parameter set at 2, from the “pracma” R package.

### Evolutionary analyses

Protein sequences used for evolutionary analyses were aligned using MAFFT version 7 with the “L-INS-i” strategy ^30^. Alignments were visualized using Jalview 2.11.1.0 ^35^. Phylogenic analyses were performed using MEGA10.1.8 software ^36^. Phylogeny trees were generated with the Maximum Likelihood statistical method, Jones-Taylor-Thornton (JTT) substitution model, uniform rates among sites, use of all sites, Nearest-Neighbor-Interchange (NNI) heuristic method, and default NJ/BioNJ initial tree. For spike protein analysis, all obtained sequences were used for alignment and phylogeny. For NSP12 analysis, all obtained ORF1ab sequences and reference SARS-CoV-2 NSP12 (YP_009725307) were used for alignment, but only ORF1ab sequences were used for phylogeny.

### Structural analyses

The crystal structure of SARS-CoV-2 spike receptor-binding domain bound with ACE2 was obtained from Protein Data Bank (PDB) with accession code 6M0J ^10^. The cryo-EM structure of the SARS-CoV-2 NSP12-NSP7-NSP8 complex bound to the template-primer RNA and the triphosphate form of remdesivir (RTP) was obtained from PDB with accession code 7BV2 ^16^. The crystal structure of SARS-CoV spike RBD bound with ACE2 was obtained from PDB with accession code 2AJF ^14^. Molecular graphics and analyses including mapping of COPA scores onto structures were performed with UCSF ChimeraX version 0.94 ^37^.

### B cell epitope analysis

FASTA sequences for reference SARS-CoV-2 structural proteins were used to predict B cell epitopes. Linear B-cell epitopes probability scores were obtained using BepiPred-2.0 ^38^. “Consensus Regions” were defined as amino acid residues with epitope scores > 0.5 and COPA scores > 8. Hypergeometric test of overlap of high COPA score (> 8) and high epitope score (> 0.5) residues was performed to determine the statistical significance of consensus regions. “Compound Regions” were identified using k-means clustering. Briefly, the R function “kmeans” was run with variable number of clusters and nstart parameter 25 on a dataset containing residue position, epitope score, and COPA score. Residues were marked as compound regions if they belonged to clusters with epitope score centers > 0.5 and COPA score centers > 8. Flagged residues that did not belong to a contiguous run of amino acids ≥ 5 residues were filtered out.

### T cell epitope analysis

FASTA sequences for reference SARS-CoV-2 structural proteins and select nonstructural proteins were used to predict T cell epitopes. Prediction of peptides binding to MHC class I and class II molecules was then performed using TepiTool ^39^ from the Immune Epitope Database (IEDB) Analysis Resource. MHC-I binder predictions were made for the “Human” host species and the 27 most frequent A & B alleles in the global population. Default settings for low number of peptides (only 9mer peptides), IEDB recommended prediction method, and predicted percentile rank cutoff ≤ 1.0 were used for peptide selection. MHC-II binder predictions were made for the “Human” host species using the “7-allele method” (median of percentile ranks from DRB1*03:01, DRB1*07:01, DRB1*15:01, DRB3*01:01, DRB3*02:02, DRB4*01:01, DRB5*01:01). Median consensus percentile rank ≤ 20.0 was used for peptide selection. Pathogenicity associated peaks within the proteins with NT-COPA scores greater than 8 were then mapped to the predicted peptides for prioritization.

### Statistical information summary

Comprehensive information on the statistical analyses used are included in various places, including the figures, figure legends and results, where the methods, significance, p-values and/or tails are described. All error bars have been defined in the figure legends or methods. Standard statistical calculations such as Spearman’s rho were performed in R with functions such as “cor”.

### Code availability

Codes used for data analysis or generation of the figures related to this study are available upon request to the corresponding author, and will be deposited to GitHub upon publication for free public access.

### Data and resource availability

All relevant processed data generated during this study are included in this article and its supplementary information files. Raw data are from various sources as described above. All data and resources related to this study are freely available upon request to the corresponding author.

## Supplementary Figure Legends

**Figure S1.**
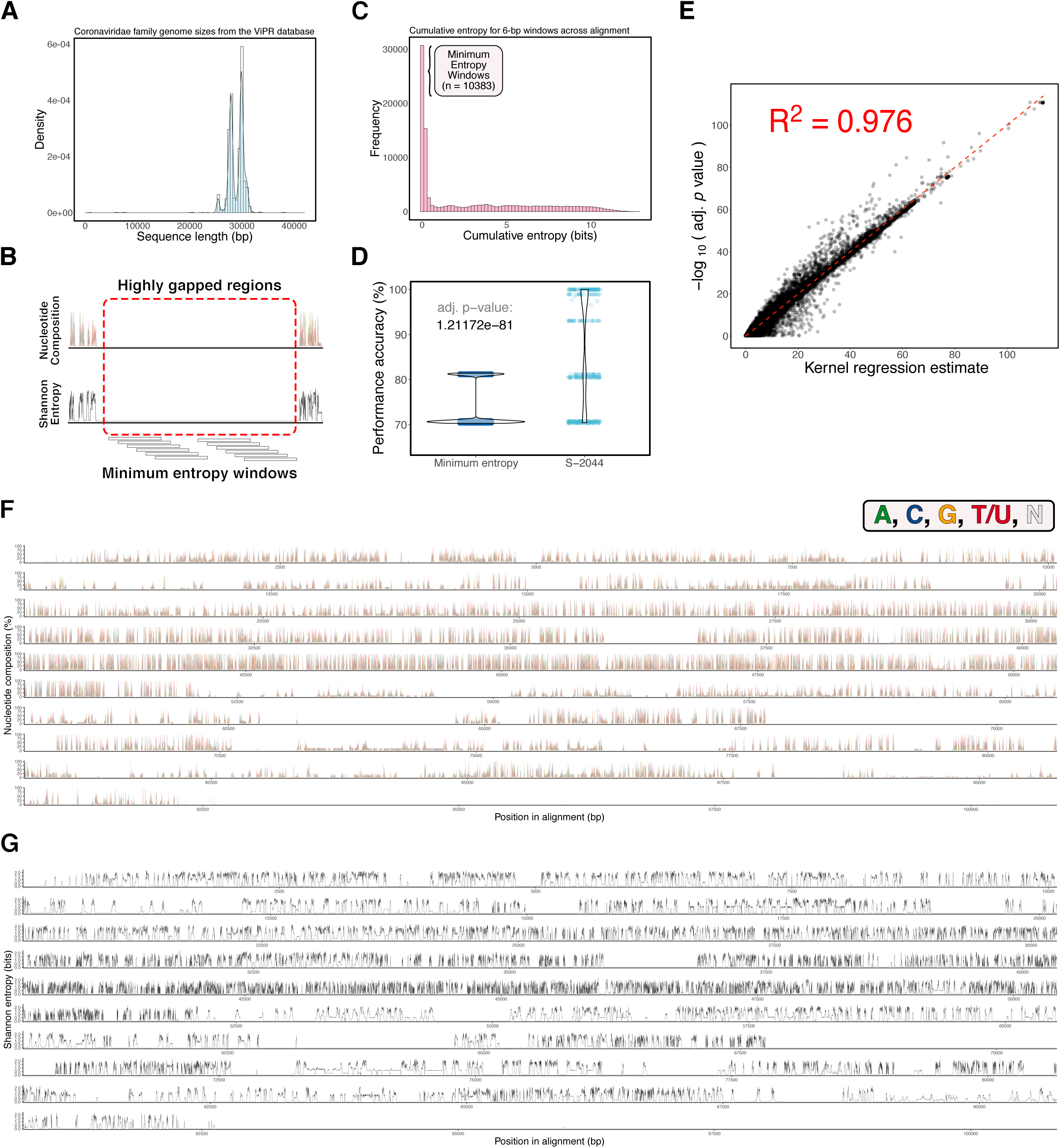
Alignment and statistical meta-model metrics. **(A)** Density histogram of total sizes in nucleotide base pair lengths for genomes. **(B)** Schematic highlighting gapped regions of genome alignment containing “minimum entropy” windows. **(C)** Histogram showing cumulative entropy scores (summed Shannon entropy values for each bp across the window) for six bp windows tiled across alignment. Of the 100,835 windows, 10,383 are classified as “minimum entropy,” corresponding to a cumulative entropy score of 0.02174422. **(D)** Example of meta-model statistical testing for position S-2044 compared to minimum entropy windows with scores learned from base ML models. P-values were obtained with two-sided Wilcoxon rank-sum test and adjusted for false discovery rate. **(E)** COPA scores plotted against kernel regression estimate. Dashed red line shows linear regression model fitted using the lm function in R, with coefficient of determination 0.976. **(F)** Trace of nucleotide composition (A, C, G, T/U, N) represented as percentage of all nucleotides shown for each position across the alignment. **(G)** Trace of Shannon entropy score calculated for each position across the alignment.

**Figure S2.**
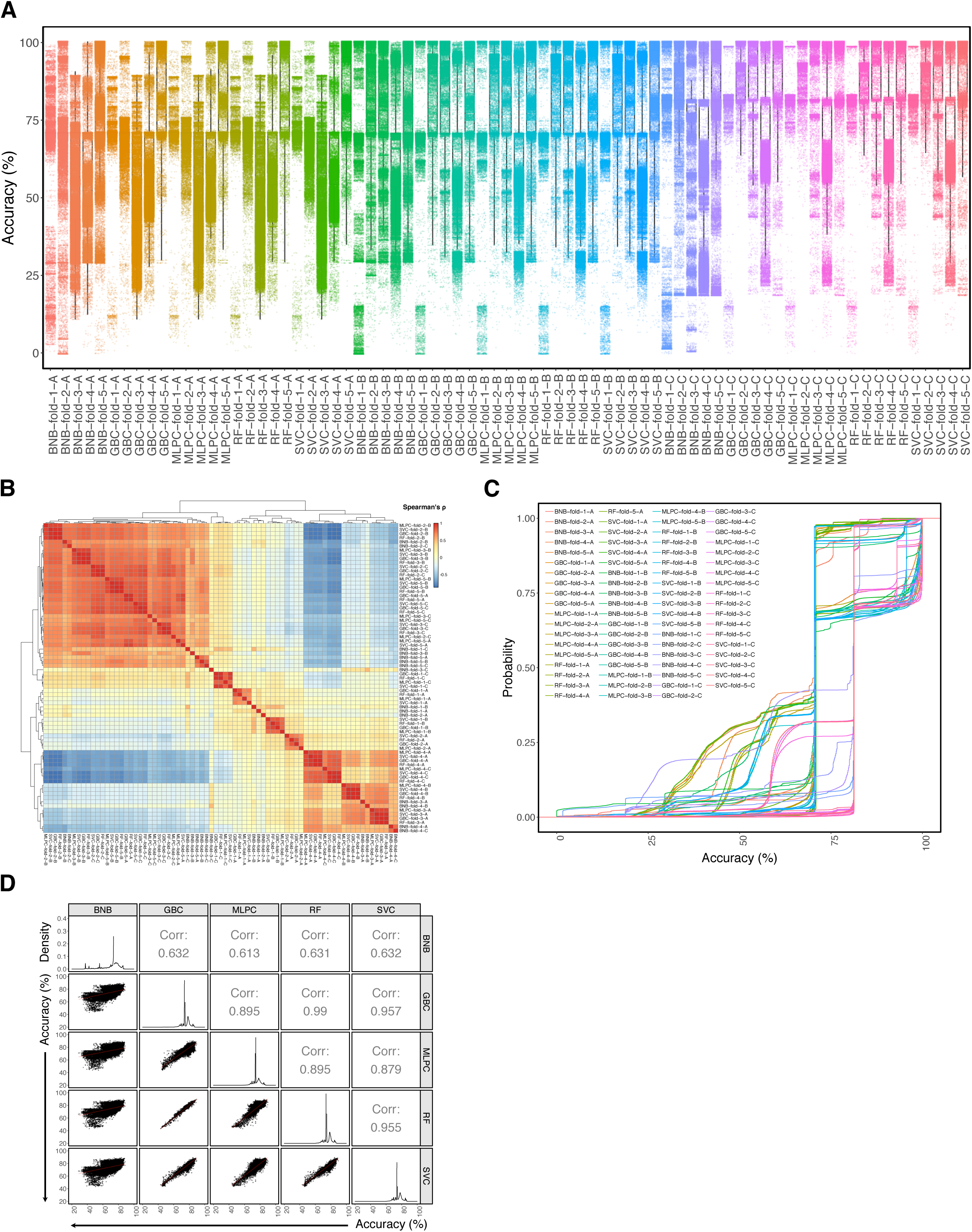
Base ML model performance metrics for standard dataset. **(A)** Scatterplots overlaid on Tukey boxplots detailing cross validation score distributions for permutations of machine learning algorithm, fold, and classification strategy (A, B, or C). **(B)** Heatmap of Spearman’s rho correlations calculated from scores for permutations of machine learning algorithm, fold, and classification strategy (A, B, or C). **(C)** Empirical cumulative distribution function of cross validation scores for permutations of machine learning algorithm, fold, and classification strategy (A, B, or C). **(D)** Pairwise comparisons plot showing correlations (Spearman’s rho) of mean window scores for different ML methods.

**Figure S3.**
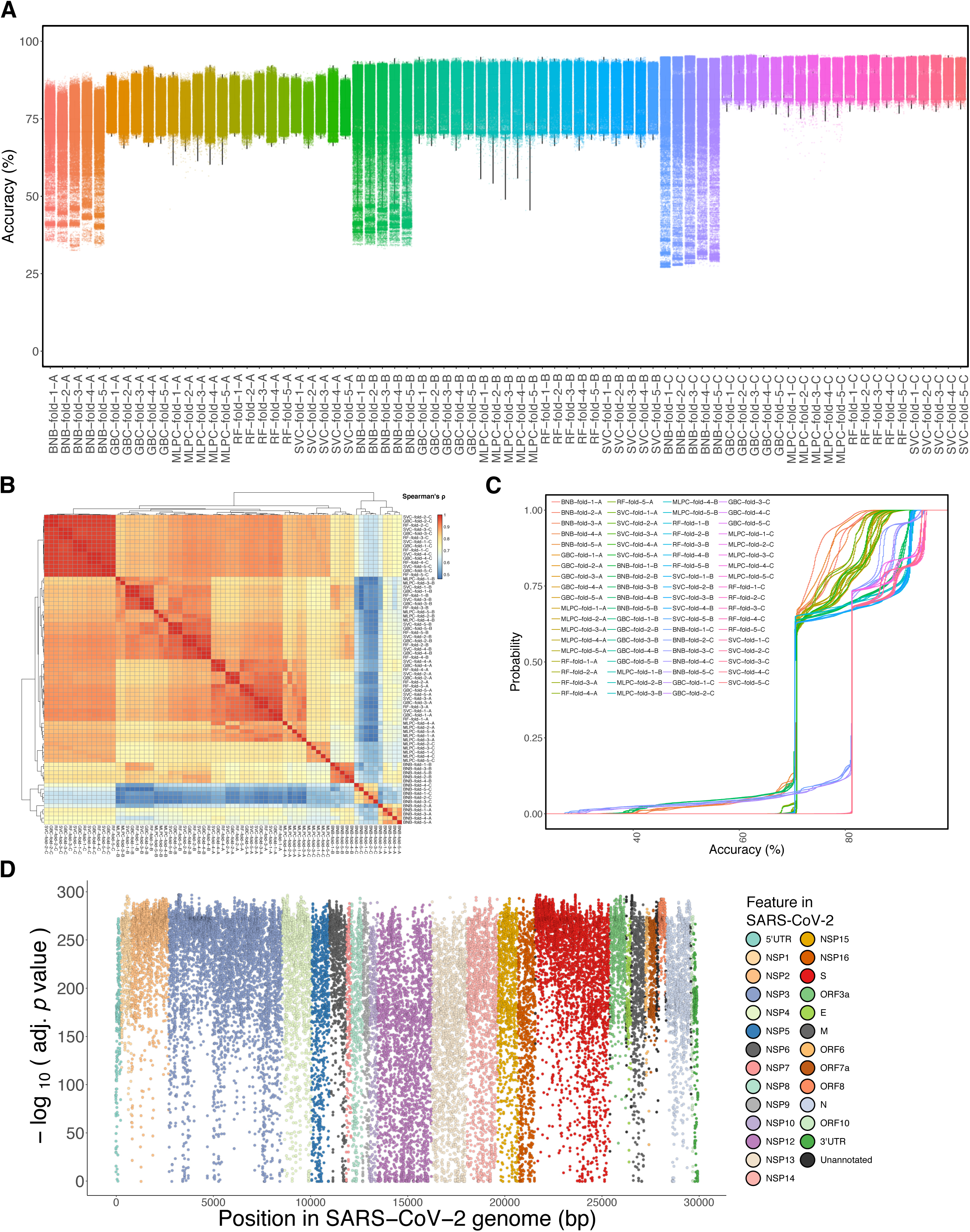
Base ML model performance metrics for shuffled dataset. **(A)** Scatterplots overlaid on Tukey boxplots detailing cross validation score distributions for permutations of machine learning algorithm, fold, and classification strategy (A, B, or C). **(B)** Heatmap of Spearman’s rho correlations calculated from scores for permutations of machine learning algorithm, fold, and classification strategy (A, B, or C). **(C)** Empirical cumulative distribution function of cross validation scores for permutations of machine learning algorithm, fold, and classification strategy (A, B, or C). **(D)** NT-COPA scores obtained by ML training on shuffled dataset for nucleotide positions across the reference SARS-CoV-2 genome.

**Figure S4.**
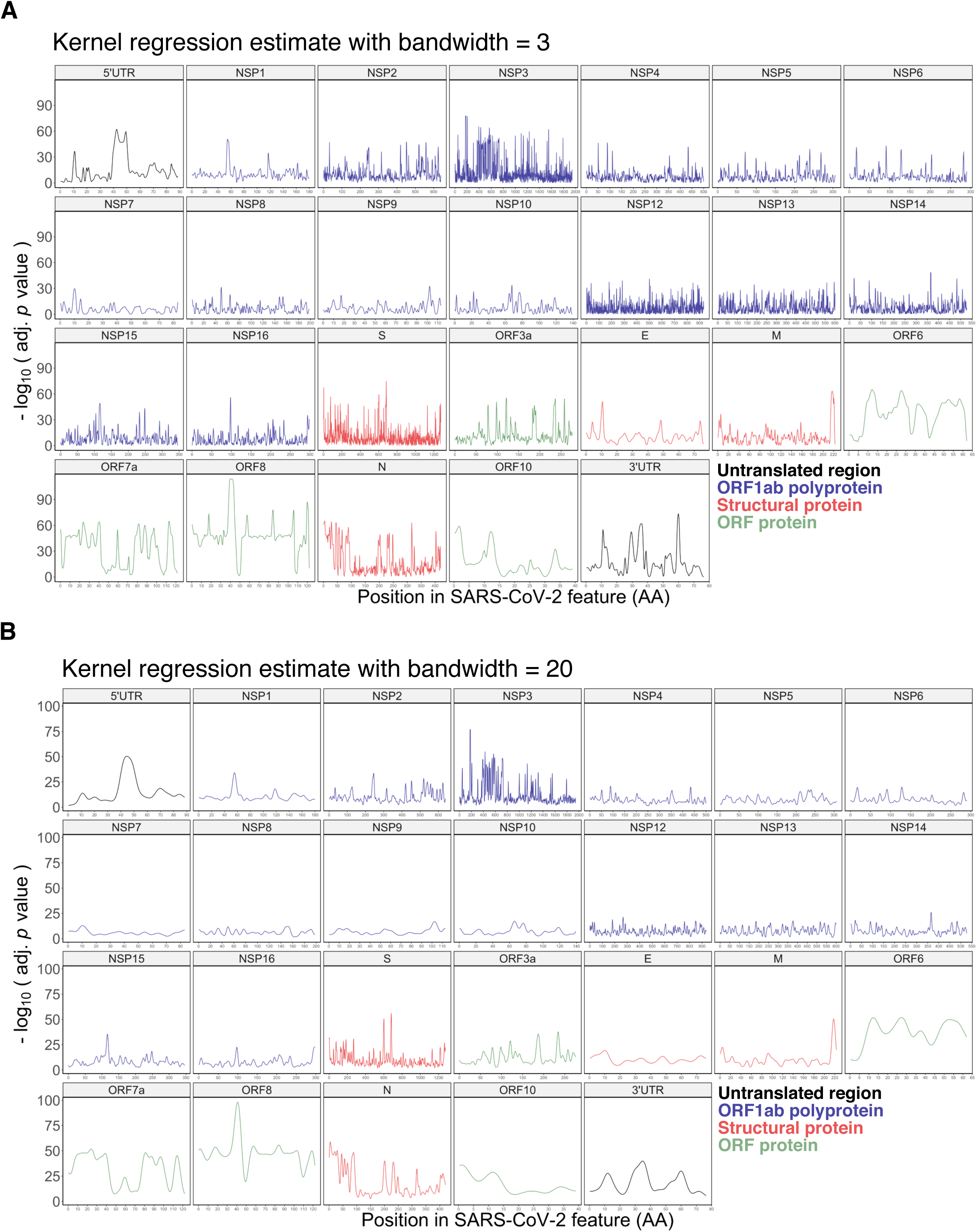
Smoothed pathogenicity signal by feature in SARS-CoV-2. **(A)** Kernel regression estimate with bandwidth = 3 obtained from COPA scores shown for individual feature in the SARS-CoV-2 genome. **(B)** Kernel regression estimate with bandwidth = 20 obtained from COPA scores shown for individual feature in the SARS-CoV-2 genome.

**Figure S5.**
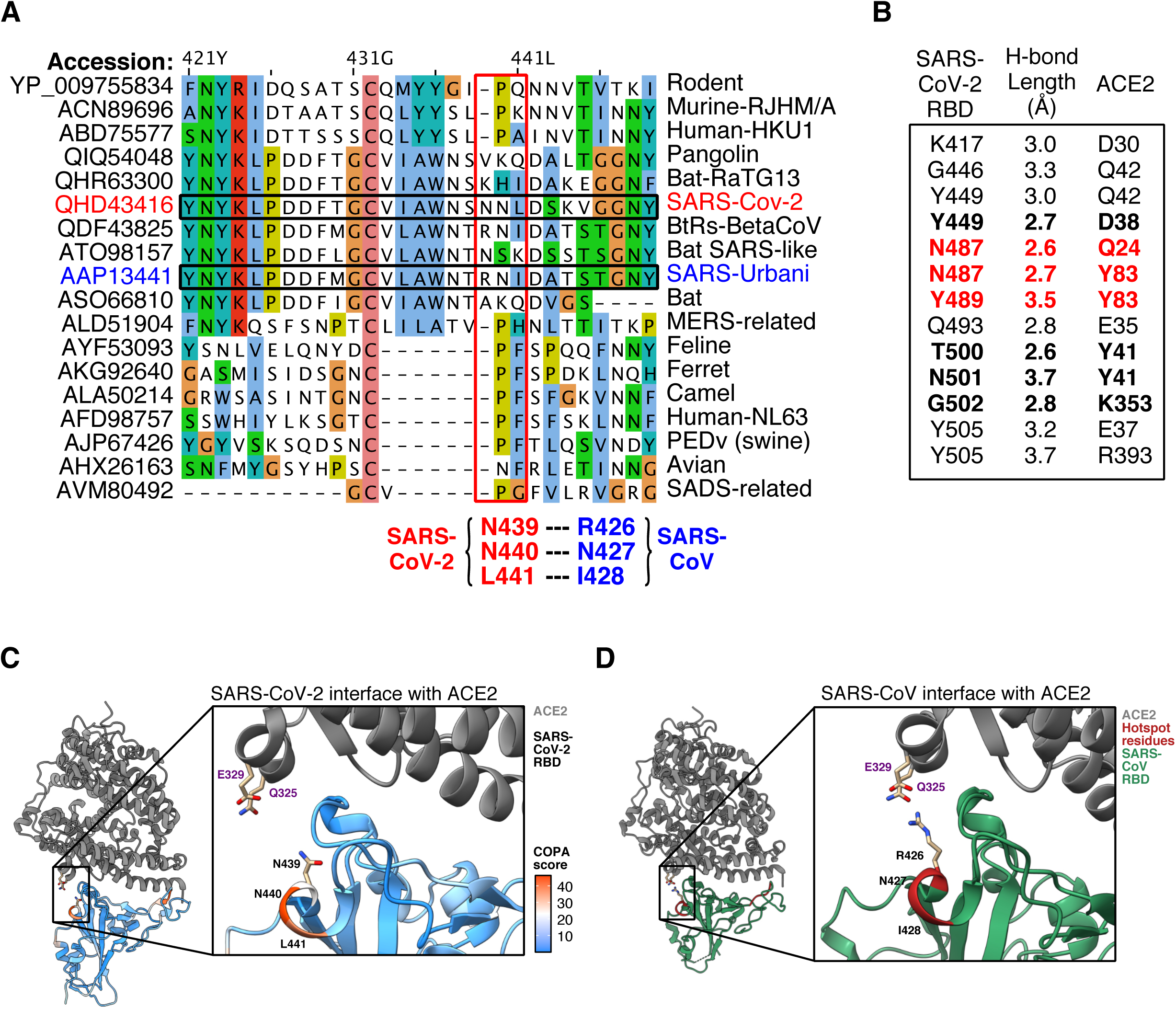
Additional evolutionary and structural analyses of NNL/RNI pathogenicity hotspot in spike protein. **(A)** Protein alignment reveals NNL hotspot for SARS-CoV-2 (RNI hotspot for SARS-CoV) has high sequence divergence from other coronaviruses across species and hosts. Residues are colored using the Clustal X color scheme. Hotspot residues for SARS-CoV-2 labelled in red, with corresponding residues for SARS-CoV labelled in blue. **(B)** Residues involved in hydrogen bonds at the SARS-CoV-2 receptor binding domain (RBD) and ACE2 interface (Lan et al., 2020). **(C)** Relative positions of SARS-CoV-2 NNL hotspot residues with ACE2 residues Q325 and E329 shown in structure of SARS-CoV-2 RBD complexed with ACE2 receptor. **(D)** Relative positions of SARS-CoV RNI hotspot residues with ACE2 residues Q325 and E329 shown in structure of SARS-CoV RBD complexed with ACE2 receptor.

**Figure S6.**
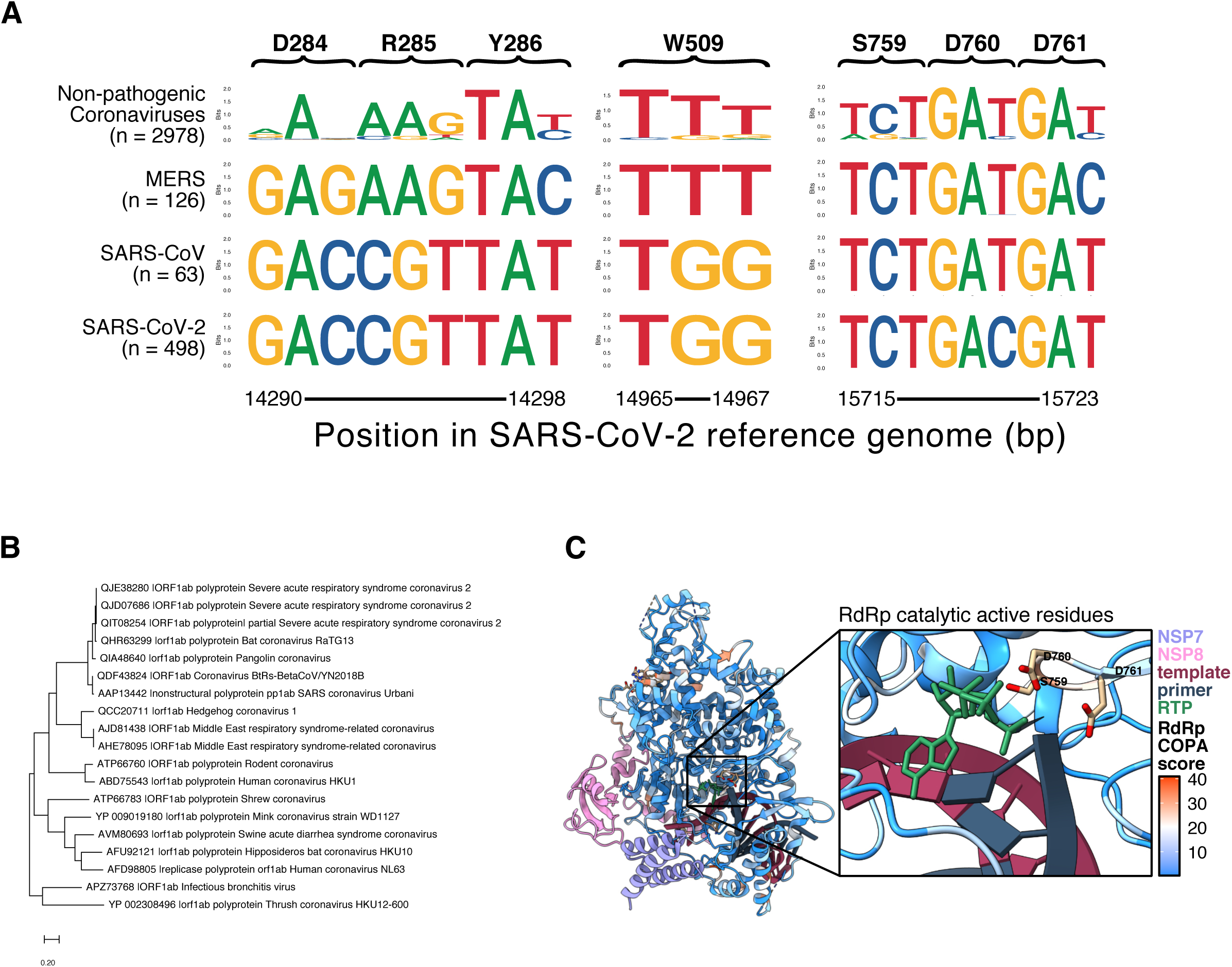
Additional evolutionary and structural analyses of key pathogenic sequences identified in SARS-CoV-2 RdRp. **(A)** Sequence logos for codons from the genome alignment used for ML training associated with the hotspot region DRY, hotspot residue W509, and catalytically active residues SDD. **(B)** Phylogeny tree for ORF1ab polypeptide sequences of coronaviruses across species and hosts. **(C)** Position and COPA scores of SARS-CoV-2 catalytically active residues SDD.

## Supplementary Datasets

### Supplementary Dataset Part A

Dataset S1. Metadata and classification strategy predictor class for Coronaviridae family genomes obtained from ViPR database.

Dataset S2. Cumulative Shannon entropy scores for six bp windows tile across alignment.

Dataset S3. Meta-model q-values, NT-COPA scores, and kernel regression estimate for each nucleotide position in SARS-CoV-2 genome with standard dataset.

Dataset S4. Meta-model q-values, NT-COPA scores, and kernel regression estimate for each nucleotide position in SARS-CoV-2 genome with shuffled dataset.

Dataset S5. Amino acid resolution COPA scores for residues in SARS-CoV-2 features.

Dataset S6. Metadata for coronavirus spike protein sequences used for evolutionary analysis.

Dataset S7. Metadata for coronavirus ORF1ab polypeptide and NSP12 protein sequences used for evolutionary analysis.

### Supplementary Dataset Part B

Dataset S8. B cell epitope scores, COPA scores, and key regions for spike protein.

Dataset S9. B cell epitope scores, COPA scores, and key regions for membrane protein.

Dataset S10. B cell epitope scores, COPA scores, and key regions for envelope protein.

Dataset S11. B cell epitope scores, COPA scores, and key regions for nucleocapsid protein.

Dataset S12. T cell epitopes and peak counts for spike protein and MHC class I.

Dataset S13. T cell epitopes and peak counts for spike protein and MHC class II.

Dataset S14. T cell epitopes and peak counts for NSP12 and MHC class I.

Dataset S15. T cell epitopes and peak counts for NSP12 and MHC class II.

Dataset S16. T cell epitopes and peak counts for nucleocapsid protein and MHC class I.

Dataset S17. T cell epitopes and peak counts for nucleocapsid protein and MHC class II.

Dataset S18. T cell epitopes and peak counts for membrane protein and MHC class I.

Dataset S19. T cell epitopes and peak counts for membrane protein and MHC class II.

Dataset S20. T cell epitopes and peak counts for envelope protein and MHC class I.

Dataset S21. T cell epitopes and peak counts for envelope protein and MHC class II.

Dataset S22. T cell epitopes and peak counts for NSP3 and MHC class I.

Dataset S23. T cell epitopes and peak counts for NSP3 and MHC class II.

Dataset S24. T cell epitopes and peak counts for NSP5 and MHC class I.

Dataset S25. T cell epitopes and peak counts for NSP5 and MHC class II.

Dataset S26. T cell epitopes and peak counts for NSP8 and MHC class I.

Dataset S27. T cell epitopes and peak counts for NSP8 and MHC class II.

Dataset S28. T cell epitopes and peak counts for NSP9 and MHC class I.

Dataset S29. T cell epitopes and peak counts for NSP9 and MHC class II.

### Supplementary Dataset Part C

Dataset S30. Classifier performance accuracy scores for permutations of machine learning algorithm, fold, and classification strategy (A, B, or C) with standard dataset.

Dataset S31. Classifier performance accuracy scores for permutations of machine learning algorithm, fold, and classification strategy (A, B, or C) with shuffled dataset.

## Notes

### Competing Interest Statement

The authors have declared no competing interest.

